# Directed evolution of a compact TranC11a system for efficient genome editing

**DOI:** 10.1101/2025.09.28.678669

**Authors:** Zixu Zhu, Quan Gao, Qiang Gao, He Jia, Zhiwei Wang, Mingyang He, Lijuan Li, Lixiao Zhang, Shengnan Li, Shuai Jin, Caixia Gao, Kevin Tianmeng Zhao

## Abstract

The recently discovered TranC systems represent programmable RNA-guided DNA endonucleases of transposon origin with compact protein sizes ideal for therapeutic delivery. However, their editing efficiency in human and plant cells is limited. Here, we evolved TranC11a and engineered its sgRNA to enhance overall editing efficiencies. TranC11a systems exhibit up to 9.2-fold higher editing activity than its parent and achieves efficiency comparable to SpCas9 across multiple human genome endogenous sites, significantly outperforming compact editors NovaIscB and enTnpB1c. TranC11a enables efficient editing of disease-relevant genes in human cells and breeding traits in maize. With its high editing activity and compact size (574 aa), TranC11a demonstrates strong potential for future *in vivo* genome editing and crop engineering.

## Main

Programmable RNA-guided DNA endonucleases represent core tools in the field of genome editing, among which CRISPR-Cas systems have already been widely used in treating genetic diseases or breeding new crops^1, 2^. Since the initial application of *Streptococcus pyogenes* Cas9 (SpCas9) for mammalian gene editing, a variety of novel nuclease systems with expanded target range or reduced size have continuously emerged^3-8^. TranC enzymes are newly identified RNA-guided DNA targeting systems, and are considered to be one of the earliest evolutionary intermediates between <u>Tran</u>sposon-associated TnpB proteins and type V <u>C</u>RISPR systems^9^. Their compact protein size (∼300-600 amino acids)^9^ highlights immense potential for use as genome editing tools, especially in clinical applications where size limitations hinder delivery. However, the editing efficiency of current TranC systems remain low in human and plant cells, limiting their practical applicability. Although other small programmable RNA-guided DNA endonucleases such as TnpB, IscB, and others have been engineered recently, their overall efficiency remains low when evaluated across multiple endogenous target sites^10-12^. Therefore, there remains a need to develop and engineer an efficient and compact nuclease for genome editing applications.

To enhance the editing performance of TranC enzymes, we integrated structural biology insights with a combination of rational design and directed evolution approaches. Using a dual-plasmid selection system, we performed saturation mutagenesis at 192 amino acid positions in eLaTranC under high and low selection pressure stringencies. Machine learning-guided mutagenesis approaches were employed to guide and predict for additional beneficial mutations. Through a series of directed evolution efforts, we obtained TranC11a, a mutant with six novel mutations (A76T, I99L, N109C, K112Q and E493V) and exhibiting over seven times enhanced editing efficiency in human cells compared to its parental eLaTranC enzyme. Furthermore, we generated multiple engineered variants of the guide RNA and found that sgC2, which is comprised of two deletions (−108 to −114 and −127 to −131) and a “C” insertion between positions −141 and −142, further increased TranC11a editing activity by 1.6-fold.

Although recent advances in compact editing systems, such as enTnpB1c^10^ and NovaIscB^11^, showed improvements in activity, their editing efficiencies generally remain lower than that of SpCas9. In this study, we found that TranC11a achieved editing efficiencies (65.8% average across eight genomic sites) superior to both enTnpB1c (17.5% average across eight genomic sites) and NovaIscB (8.8% average across eight genomic sites), and even comparable to or superior when compared with SpCas9 (51.9% average across eight genomic sites). Importantly, we selected the most proximal protospacer for each site based on PAM (or TAM) availability to limit any potential bias due to local chromatin states. Finally, we preliminarily validated the application potential of TranC11a in both plant protoplasts and human cells, for plant breeding and gene therapy, respectively. This study not only enhances the editing performance of the TranC system but also provides reliable tools for its practical application in plant and mammalian genome editing.

## Results

### Structure- and AI-guided directed evolution of TranC enzymes

Due to TranC’s low editing activity, we embarked on a structure-guided directed evolution campaign to increase TranC mammalian editing efficiencies (**Fig. 1b**). Given that most residues in LaTranC (TranC enzyme from *Lachnospiraceae sp*.) had already been subjected to five rounds of arginine-scanning mutagenesis^9^, we opted to use site-saturation mutagenesis to generate a comprehensive library of variants which could then be combined to design for variants with activity increases. Rather than performing saturation mutagenesis at all amino acid residues, we focused on 192 amino acid positions based on different key insights: (1) PAM-interacting residues and (2) nucleic acid-proximal residues based on a predicted structure of LaTranC–sgRNA–DNA by AlphaFold3^13^ and (3) high sequence variability residues defined as positions upon which a substitution would display high variance in editing efficiency based on a machine learning model developed using previous mutants of LaTranC^9^. Additionally, we evaluated (4) specific mutations proposed by the machine learning model when trained with all previously developed mutants of eLaTranC^9^ (**Fig. 1b, Methods**).

**Fig. 1:**
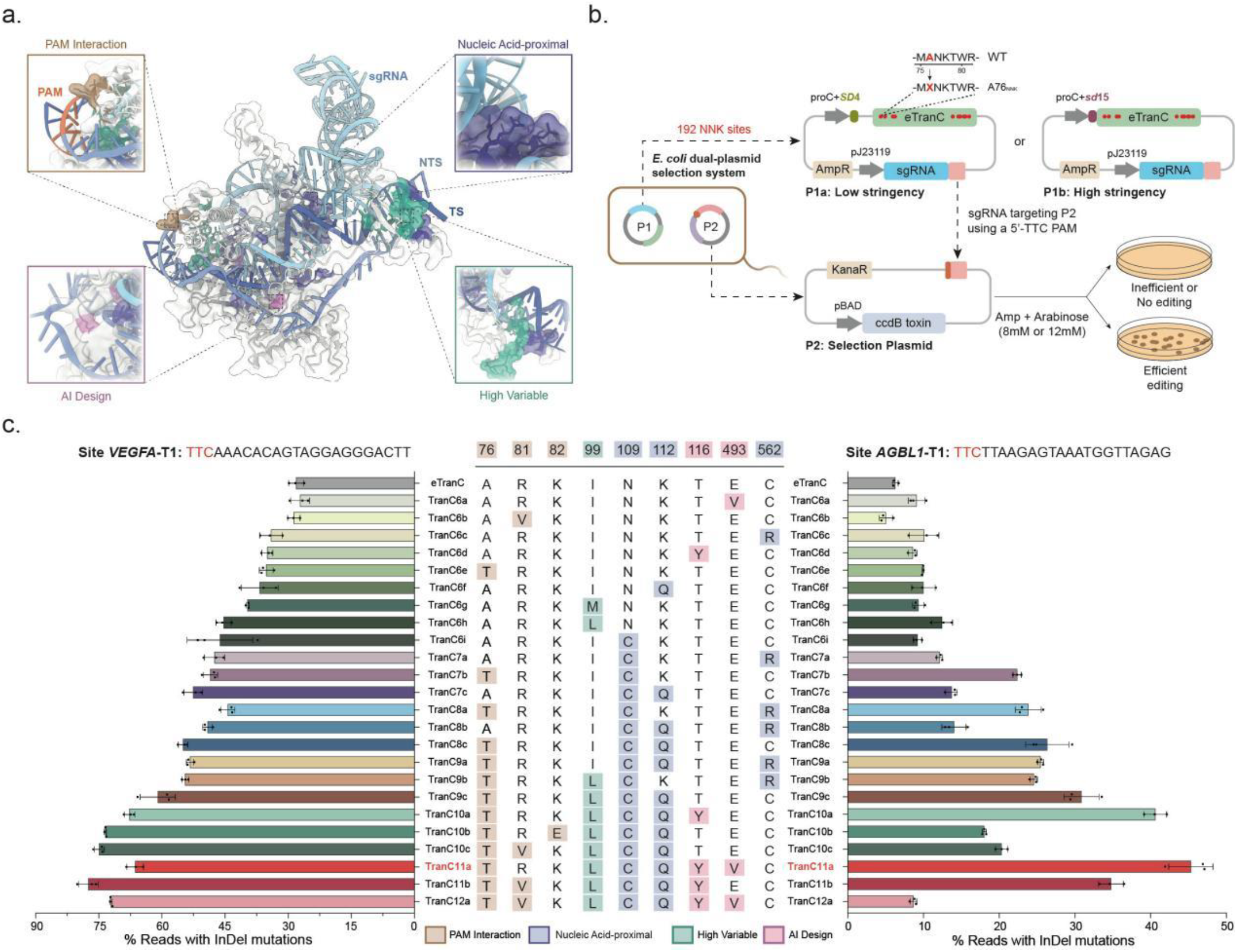
Directed evolution of TranC11a protein. **a**, AlphaFold3-predicted structure of eLaTranC-sgRNA-target DNA ternary complex (Site *VEGFA*- T1). The amino acid residues selected for evolution are colored in different colors. Brown, PAM-interacting residues; purple, nucleic acid-proximal residues; pink, AI-designed mutants; green, high variable residues; light grey, eLaTranC; light blue, sgRNA; blue, NTS; Dark blue, TS. **b**, Schematic diagram of the *E. coli* dual-plasmid selection system. Two different screening stringencies were used during the evolution (P1a and P1b). **c**, Editing efficiencies of eLaTranC and 24 representative variants throughout the evolution campaign in HEK293T cells at Site *VEGFA*-T1 and Site *AGBL1*-T1. Values and error bars represent the means and s.d., respectively, for three independent biological replicates.

To select for TranC variants with enhanced nuclease activities, we established a positive selection system in *E. coli* based on the cytotoxic *ccdB* gene (**Fig. 1b** and **Supplementary Fig. 1a**). In this system, we encoded the expression of a TranC enzyme and its guided RNA on a P1 plasmid. The guide RNA is constitutively expressed under a J23119 promoter while the TranC enzyme is expressed by either a proC promoter coupled with a *SD4* RBS for low stringency selections (P1a), or a proC promoter coupled with a *sd15* RBS for high stringency selections (P1b). On the selection plasmid, P2, a *ccdB* gene is expressed by an arabinose-inducible pBAD promoter (**Fig. 1b**). We identified a 24-nt target site following a 5’-TTC PAM to serve as the target for the TranC enzyme to cleave on P2 (**Supplementary Table 1**); successful cleavage of P2 by active TranC variants would promote bacterial survival upon induction with arabinose during selection.

Initial validations showed that the expression of a catalytically inactive TranC variant (D253A) failed to rescue bacterial growth upon arabinose induction, whereas a functional eLaTranC permitted survival, confirming the design of the selection circuit (**Supplementary Fig. 1a**). We next optimized combinations of post-electroporation recovery times and arabinose concentrations to ensure proper selection strengths for both stringency levels (**Supplementary Fig. 1b, Methods**). Under a low-stringency condition, eLaTranC expressed using P1a with a 10-minute recovery time post-electroporation and 8 mM arabinose induction for selection showed minimal background survival (**Supplementary Fig. 1b**). Therefore, we elected to use this condition for the first round of library screening. Next, we designed the high-stringency selection to express eLaTranC using P1b coupled with a 20-minute recovery time post-electroporation followed by 12 mM arabinose induction for survival, which results in complete elimination of any background colony growth (**Fig. 1b, Supplementary Fig. 1b**).

To develop TranC saturation mutagenesis libraries, we split the targeted 192 selected amino acid residues into 24 subpools, with each encompassing a universal reverse primer with unique forward primers to generate NNK-libraries at each selected amino acid residues. We next combined all 24 subpools into 8 combined pools based on amino acid distance (60 amino acids) for the selection. Using the P1a and P1b screening conditions described above, we subjected the saturation mutagenesis library to both conditions. In total, we obtained over 50,000 colonies and analyzed 1,129 using Sanger or amplicon sequencing (**Supplementary Table 1**, **Methods**).

A total of 147 mutants, which were selected based on their occurrence frequency (**Supplementary Table 1**), were chosen to be evaluated in eukaryotic cells. We cloned 147 single mutants derived from the selection and 37 AI-designed single mutants (**Methods**) into mammalian cell-expressing vectors to evaluate their editing activity in HEK293T cells across two endogenous genomic sites (Sites *VEGFA*-T1 and *AGBL1*-T1) (**Supplementary Table 1**). Among the 147 mutants identified from the bacterial selection, 13 showed improved efficiencies over eLaTranC at both sites, while 6 out of the 37 AI-designed variants exhibited enhanced activity (**Supplementary Fig. 1c, d**). eLaTranC is in itself comprised of five mutations (S137R, P148R, D150R, K315R and A369R) on top of the wildtype LaTranC enzyme, so future naming mechanisms focus on the total number of mutations introduced including the five in eLaTranC. The most efficient variant at Site *VEGFA*-T1 is TranC6i (N109C, 46.2%), representing a 1.6-fold improvement over eLaTranC, while the most efficient variant at Site *AGBL1*-T1 is TranC6h (I99L, 12.5%), representing a 1.5-fold increase over eLaTranC (**Fig. 1c**).

We subsequently generated combinatorial mutants by introducing C562R, A76T and K112Q into TranC6i resulting in TranC7a (N109C+C562R), TranC7b (N109C+A76T), and TranC7c (N109C+K112Q) (**Fig. 1c**). All three combinational variants exhibited higher editing efficiencies than TranC6i at both sites. TranC7c showed the highest activity at Site *VEGFA*-T1 (52.6%, 1.6-fold of eLaTranC), while TranC7b was most efficient at Site *AGBL1*-T1 (22.4%, 3.6-fold of eLaTranC) (**Fig. 1c**).

Using TranC7b and TranC7c as templates, we next combined and shuffled in other mutations to yield TranC8a (A76T+N109C+C562R), TranC8b (A76T+N109C+C562R), and TranC8c (A76T+N109C+K112Q). TranC8c demonstrated the highest editing efficiency at both sites, resulting in 55.0% and 26.4% at Site *VEGFA*-T1 and Site *AGBL1*-T1, respectively (2.0-fold and 4.2-fold over eLaTranC) (**Fig. 1c**). Using TranC8c as a template, we introduced additional point mutations to generate 9-mutation variants, among which TranC9c (A76T+N109C+K112Q+I99L) showed the highest editing efficiency (60.9% at *VEGFA*-T1 and 30.9% at *AGBL1*-T1) (**Fig. 1c**). Subsequent rounds of mutation combination involved the construction and testing of 10-, 11-, and 12-mutation variants. We found that the 12-mutation variants exhibited substantial decreases in overall editing efficiency, while TranC11b and TranC11a emerged as the top performers at Site *VEGFA*-T1 (77.5%, 2.8-fold over eLaTranC) and Site *AGBL1*-T1 (45.3%, 7.2-fold over eLaTranC), respectively (**Fig. 1c**). Because TranC11a exhibited superior activity at the more refractory site, *AGBL1*-T1, we selected TranC11a (A76T+I99L+N109C+K112Q+E493V) as the final variant for subsequent evaluations (**Fig. 1c**).

### TranC sgRNA engineering

Following the evolution of TranC11a, we next wondered if the sgRNA component of TranC systems could be further enhanced to promote superior editing activity. We generated a series of rationally designed sgRNAs to enhance and stabilize the RNA’s secondary structure. Based on the LaTranC cryo-EM structure^9^, we designed modifications at the Stem2, Stem3, and triplex regions of the sgRNA (**Fig. 2a**). For Stem2 and Stem3, we designed four variants each (sg1–sg8) to correct for the sub-optimal base pairing at these regions. An additional variant, sg9, was engineered to introduce complementarity between the linker region connecting Stem2 and Stem3 to a segment of Stem1 (**Fig. 2a**).

**Fig. 2:**
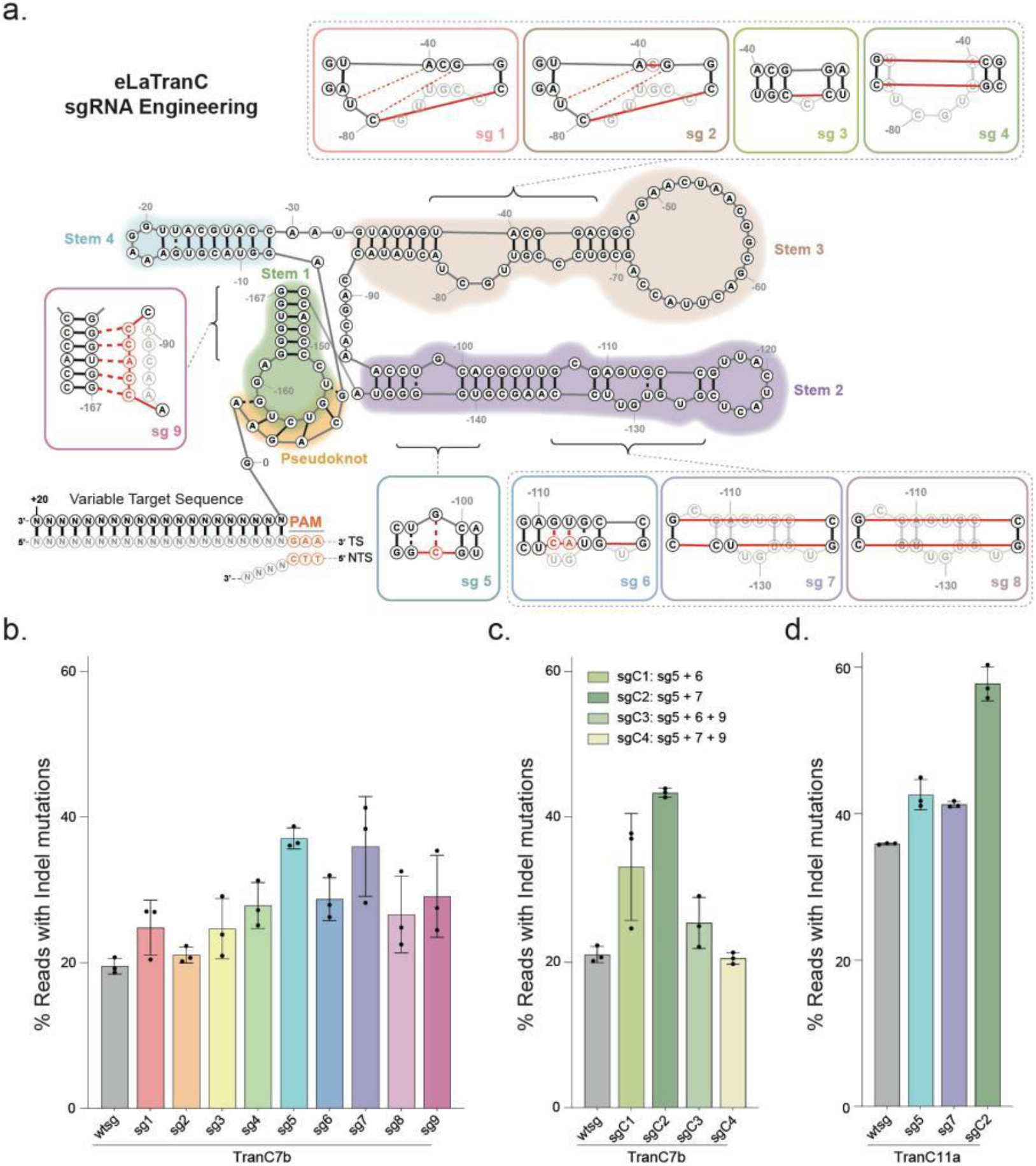
Structure-guided sgRNA engineering. **a**, Schematic diagram of the secondary structure of the eLaTranC sgRNA. Different regions of the sgRNA are colored with different background colors, and the engineering methods and locations of the nine sgRNA modifications are displayed. **b**-**c**, Editing efficiencies of TranC7b using 9 sgRNA single modifications (**b**), and 4 sgRNA combinational variants (**c**) in HEK293T cells. Values and error bars represent the means and s.d., respectively, for three independent biological replicates. **d**, Editing efficiencies of TranC11a using three best sgRNA variants compared to the wtsg. Values and error bars represent the means and s.d., respectively, for three independent biological replicates.

We first evaluated the editing activity of all nine sgRNA variants with TranC7b at Site *AGBL*-T1. TranC7b was selected as an intermediate enzyme which could better expose any differences in editing efficiency. Surprisingly, all nine variants enhanced editing efficiency, with sg5, sg7, and sg9 showing the highest activity at Site *AGBL*-T1 (**Fig. 2b**). We then combined these three modifications in different combinations to generate four composite variants, sgC1-sgC4, and evaluated their activity using TranC7b (**Fig. 2c**). Among these, sgC2—which combines the mutations from sg5 and sg7—yielded the highest editing efficiency at Site *AGBL*-T1 (**Fig. 2c**, 43.3%, 2.1-fold improvement over the wild-type sgRNA).

We next assessed the compatibility of the optimal protein variant, TranC11a, with sgC2. We compared the editing efficiencies of TranC11a with either sg5 or sg7 alone and with sgC2; as expected, we found that sgC2 (157-nt scaffold) achieved the highest editing efficiency when paired with TranC11a (**Fig. 2d**, 57.7%, representing a 1.6-fold increase over wild-type sgRNA). Consequently, we employed TranC11a in combination with sgC2 for all subsequent experiments.

### Characterization of TranC11a activity in mammalian cells

We next evaluated the editing performance of the optimized TranC11a system in comparison with other widely used genome editing enzymes. In addition to SpCas9, we compared the editing efficiency of TranC11a with two recently reported compact editors, enTnpB1c^10^ and NovaIscB^11^. Among these three compact editors, enTnpB1c has the smallest protein size (407 aa) but the largest guide RNA (230 nt). TranC11a has a slightly larger protein (574 aa) than enTnpB1c but the smallest guide RNA (157 nt). NovaIscB is the largest in protein size (613 aa), with an intermediate guide RNA length (166 nt) (**Fig. 3a**).

**Fig. 3:**
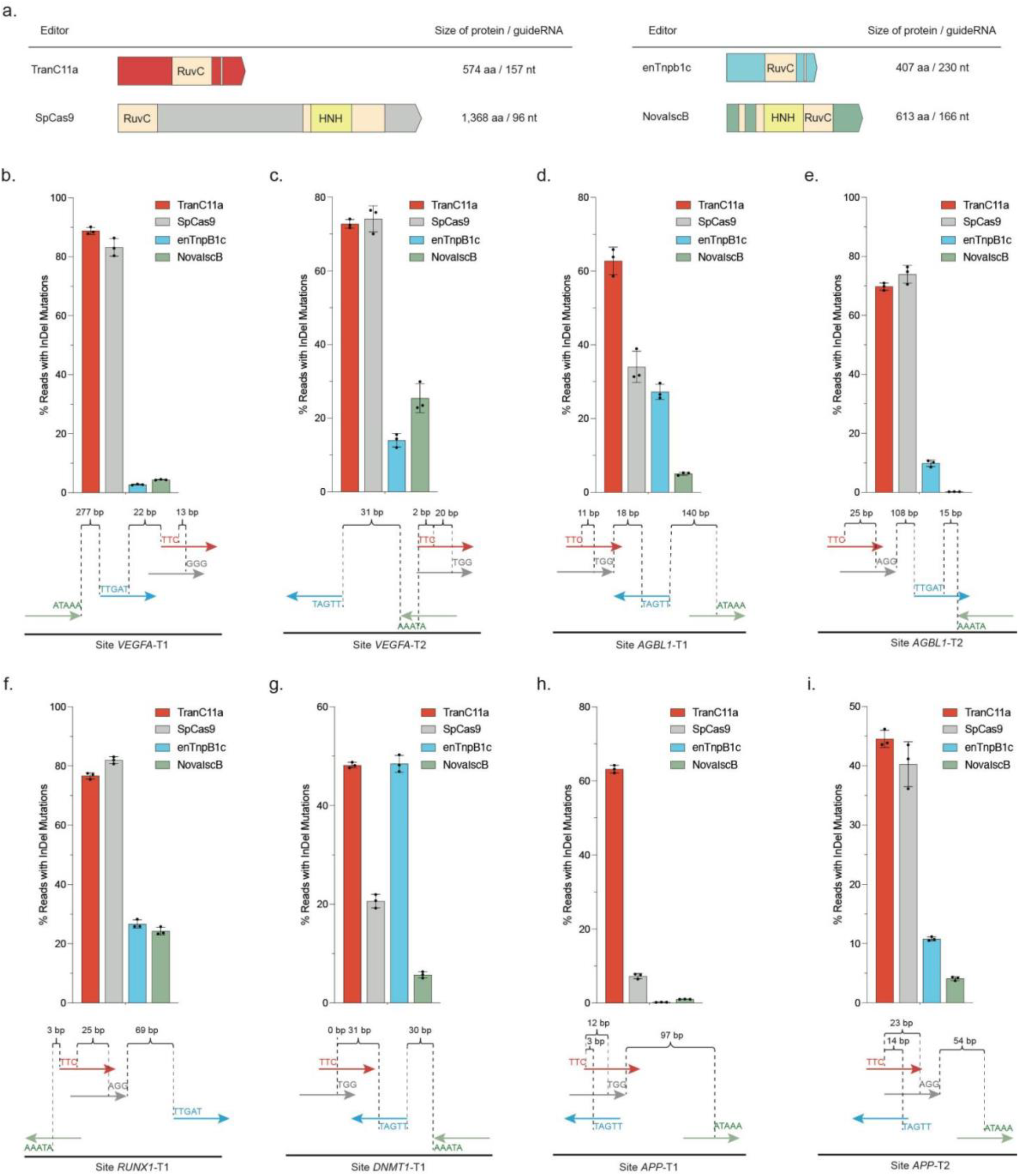
Characterization of TranC11a activity in mammalian cells. **a**, Schematic diagrams of TranC11a and three tested editors. The sizes of the protein and the corresponding guide RNA scaffold are presented. Light yellow, RuvC domain; Yellow, HNH domain. **b**-**i**, Editing efficiencies of TranC11a, SpCas9, enTnpB1c and NovaIscB at eight endogenous sites in HEK293T cells (**b**, Site *VEGFA*-T1; **c**, Site *VEGFA*-T2; **d**, Site *AGBL1*-T1; **e**, Site *AGBL1*-T2; **f**, Site *RUNX1*-T1; **g**, Site *DNMT*-T1; **h**, Site *APP*-T1; **i**, Site *APP*-T2). Target site schematic diagrams of all four editors at each genomic locus are represented by red (TranC11a, 5’-TTC), gray (SpCas9, 3’-NGG), blue (enTnpB1c, 5’-TTGAT), and light green (NovaIscB, 3’-ATAAA); the distance shown is the position from the first or last position of one PAM or TAM to the next closest protein’s PAM or TAM. Values and error bars represent the means and s.d., respectively, for three independent biological replicates.

To enable a fair comparison of editing efficiencies, we selected eight genomic target sites in HEK293T cells, each containing cognate TAM (or PAM) sequences recognized by the four editors in close proximity. Importantly, we elected to select the closest positioned protospacer for each enzyme dependent on the PAM or TAM positioning to minimize any potential chromatin effects on editing efficiency (**Fig. 3b–i**). Due to TranC11a exhibiting a relatively simple 5’-TTC PAM, most TranC11a spacers overlapped with the SpCas9 target site. TranC11a exhibited activity comparable to SpCas9 at most sites (average 65.8% vs 51.9% across eight sites) and showed on average 3.8-fold and 7.5-fold higher editing than enTnpB1c (average 17.5% across eight sites) and NovaIscB (average 8.8% across eight sites), respectively (**Figs. 3b–i**). The editing activity of enTnpB1c and NovaIscB could be lower in this case as their TAMs are complex in nature (5’-TTGAT-spacer and spacer-ATAAA-3’, respectively) so the closest positioned spacer with a respective TAM may be not optimized. However, due to their close proximity, this positioning would allow for activity comparisons while minimizing any potential effects of local chromatin architecture on editing efficiencies.

### Application of TranC11a to therapeutic targets and plant breeding

To evaluate the utility of TranC11a across translational applications, we assessed its efficacy at editing disease-relevant loci in human cells (**Fig. 4a**) and in molecular breeding applications in plant cells (**Fig. 4b**). Knocking out *B2M* minimize graft-versus-host disease and host immune rejection is a key strategy for developing ready-to-use allogeneic beta cells for treating diabetes^14^. Indeed, recent first-in-human data shows promise that such an approach would be feasible moving forward for developing novel therapeutics^15^. TranC11a mediated editing efficiencies of up to 86.3% in *B2M* across three different target sites (**Fig. 4c**).

**Fig. 4:**
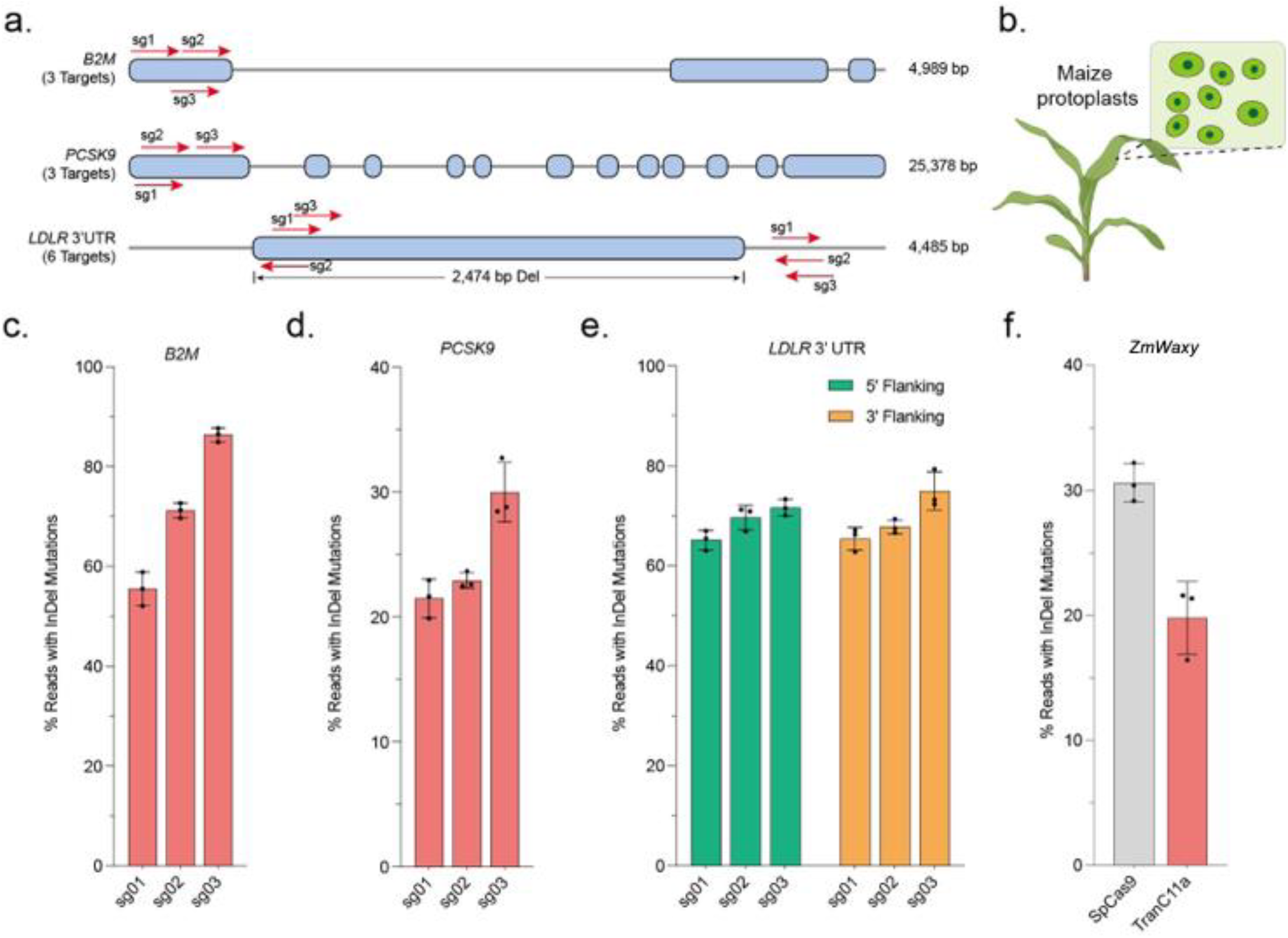
TranC11a is an efficient tool for mammalian and plant genome editing. **a**, Gene diagrams of *B2M, PSCK9*, 3’UTR regions of *LDLR*. Exons are shown as light blue boxes. Short red lines demarcate the positions of the chosen protospacers. Arrowheads indicate the 5’→3’ direction. **b**, Schematic diagram of maize protoplasts. **c**-**e**, Editing efficiencies of TranC11a at *B2M* (**c**), *PCSK9* (**d**), 5’ and 3’ flanking sequence of *LDLR* (**e**) in HEK293T cells. Values and error bars represent the means and s.d., respectively, for three independent biological replicates. **f**, Editing efficiencies of TranC11a and SpCas9 at *ZmWaxy*. Values and error bars represent the means and s.d., respectively, for three independent biological replicates.

Next, we turned towards hypercholestermia and sought to evaluate the potential of editing *PCSK9* and *LDLR*, two key proteins implicated in high cholesterol. We first tested the editing activity of TranC11a at three targets of *PCSK9*^16^ and found the highest editing efficiency reaching 30.0% (**Fig. 4d**). In parallel, recent studies have identified a naturally occurring 2.5-kb deletion within the 3’ UTR of the *LDLR* gene in an Icelandish population to be correlated with low LDL-C level (13-72 mg/dL compared to 105-120 mg/dL in normal people)^17^. To recapitulate this deletion genotype, we targeted the flanking sequences of this 2.3-kb deletion with TranC11a, achieving efficiencies of 71.7% (5’-Flanking) and 75.0% (3’-Flanking) (**Fig. 4e**).

To assess the utility of TranC11a in plants, we delivered plasmids encoding TranC11a into maize protoplasts to verify its activity in plant cells (**Fig. 4b**). Knockout of the *Waxy* gene in maize breeds for specialty varieties with altered starch quality^19^. We evaluated editing *ZmWaxy* in maize protoplasts using either TranC11a or SpCas9. The editing efficiencies of TranC11a and SpCas9 on *ZmWaxy* were 19.8% and 12.9%, respectively (**Fig. 4f**). This highlights TranC11a as a feasible enzyme to use in plant breeding applications. Future efforts will continue to evaluate suitable conditions that enhance editing efficiencies across different plant species.

## Discussion

In this study, we evolved and engineered a highly efficient and compact (574 aa) TranC-based editor, termed TranC11a. Through engineering both the protein and guide RNA components, TranC11a exhibits an up to 9.2-fold improvement in editing activity compared to eLaTranC^9^. TranC11a contains six novel mutations: A76T (PAM-interacting domain), I99L (high variable region), N109C, K112Q, C562R (all three from the nucleic acid-proximal region), T116Y and E493V (both proposed by machine learning models). Concurrently, we engineered the sgRNA scaffold by optimizing non-canonical base pairs in Stem2 and Stem3 and redesigning an inter-stem linker. A combination of these modifications resulted in sgC2 (156 nt scaffold), which further enhanced TranC11a activity.

When evaluating TranC11a across eight endogenous sites in the human genome, we found that TranC11a achieved editing efficiencies comparable to that of SpCas9. Moreover, TranC11a consistently outperformed the other compact editors NovaIscB and enTnpB1c across all tested eight target sites, positioning it as a superior and highly potent compact editor with broad utility in genome engineering. The high editing efficiency achieved by TranC11a at several therapeutic targets (*B2M, LDLR, PCSK9*) establishes proof of concept for its future application in disease treatment. Efficient editing of the agronomical-related *ZmWaxy* in maize by TranC11a demonstrates its utility as a versatile tool for molecular breeding. Because different plant cells are cultured under different conditions, additional optimizations are necessary to ensure efficient editing across different species. Future structural analyses of TranC11a may elucidate key mechanisms in how the novel mutations obtained during directed evolution aided in the enhancement of overall genome editing activity enhancement.

## Online Methods

### Plasmid construction

All PCR reactions were performed using 2× Phanta Flash Master Mix (Vazyme Biotech) unless otherwise specified. All plasmid constructions were performed using Uniclone One Step Seamless Cloning Kit (Genesand). For *E. coli* screening experiment, P1 plasmid is modified from the pCandidate vector^9^. Specifically, the eLaTranC protein-coding sequence derived from pCMV-eLaTranC^9^, the LaTranC sgRNA sequence from the phU6-sgRNA vector^9^ and the commercially synthesized proC+*SD* sequence (**Supplementary Table 1**) will be amplified and ligated into the pCandidate plasmid. P2 plasmid was modified from p11-LacY-wtx1^18^. Specifically, the KanR expression frame sequence and GFP-T1 sequence (**Supplementary Table 1**) with a 5’-TTC PAM derived from pCandidate-LaTranC were amplified and ligated into the p11-LacY-wtx1^20^ vector respectively. For the TranC mutants tested in HEK239T cells, mutations were introduced at the target amino acid sites using the reverse PCR method based on the pCMV-eLaTranC backbone^9^. For NovaIscB and enTnpB1c test in HEK239T cells, protein and sgRNA sequence were commercially synthesized and cloned into a pCMV- and phU6-backbone, respectively. Plasmids for *E. coli* screening experiment and HEK293T cell line transfection were extracted and purified using EndoFree Plasmid Kits (Qiagen) or *EasyPure*® HiPure Plasmid MiniPrep Kit (TransGene Biotech). For plasmids test in plant protoplasts, TranC11a sequence from pCMV vector were amplified and ligated into a pUBI backbone or a p35s backbone^21^. TranC11a sgC2 sequence were amplified and ligated into a p35Sen&U6 backbone. Plasmids for plant protoplasts were extracted and purified using EndoFree Plasmid Kits (GeneSand). Primers used for amplicon sequencing were synthesized by the Beijing Tsingke Biotech and are listed in **Supplementary Table 1**.

### Cell culture and transfection

HEK293T cells are cultured in DMEM (Gibco) supplemented with 10% (vol/vol) FBS (Gibco) and 1% (vol/vol) penicillin–streptomycin (Gibco). All the cells were routinely tested for Mycoplasma contamination with a mycoplasma detection kit (TransGen Biotech). For cells transfection, 6 × 10^4^ cells per well were seeded into 48-well poly-D-lysine-coated plates (Corning) in the absence of antibiotic. After 16–24 h, 200 ng TranC protein expressing plasmid together with 50 ng sgRNA expressing plasmid was transfected into the cells. After 48 h, cells were collected for DNA extraction.

### Genomic DNA isolation from mammalian cell culture

Genomic DNA extraction was performed by the addition of 100 μl freshly prepared lysis buffer (10 mM Tris–HCl, pH8.0), 0.05% SDS and 25 μg ml^−1^ proteinase K (Thermo Fisher Scientific) directly into the 48-well culture plate after cells were washed once with 1× Dulbecco’s PBS (Thermo Fisher Scientific). The mixture was incubated at 37 °C for 60 min and then treated at 80 °C in the thermocycler for 20 min.

### Deep sequencing and data analysis

Two rounds of PCR were used to amplify a DNA fragment containing the target site and perform subsequent sequencing. In the 1^st^ round of PCR, the target region was amplified from the genomic DNA with site-specific primers using the DNA template. In the 2^nd^ round PCR, both forward and reverse barcodes were added to the ends of the PCR products for library construction. Equal amounts of PCR product were pooled and purified with a FastPure Gel DNA Extraction Mini Kit (Vazyme Biotech. Inc.) and quantified with a Qubit 4 (Thermo Fisher Scientific). The purified products were sequenced using the BGI G99 or Illumina Miseq platforms, and the sequences around the target regions were examined to analyze the editing outcomes. Sequence of all the targets, NGS primers and the corresponding amplicons are listed in **Supplementary Table 1**.

### Selection of TranC sites for saturation mutagenesis

We constructed a rational protein-evolution design pipeline for the LaTranC protein as follows: first, we used AlphaFold3^13^ to predict high-confidence three-dimensional models of LaTranC in complex with both RNA and DNA, and imported these complexes into PyMOL (v3.1) for atomic-level spatial analysis; residues whose any heavy atom lies within 4.0 Å of any atom of nucleic acid (DNA or RNA) in the corresponding complex were annotated as nucleic acid-proximal, with those in the DNA complex termed “DNA-proximal residues” and those adjacent to the protospacer adjacent motif (PAM) region of DNA—specifically any residue within 4.0 Å of any atom of the PAM-containing DNA segment—termed “PAM-proximal residues”. In parallel, to identify mutation sensitive hotspots for enhancing editing efficiency, we compiled arginine-scaning data and single-point mutation data for LaTranC variants, measuring editing efficiencies of each mutant relative to wild-type; using these datasets we trained an AI-guided evolutionary model to map the sequence-function landscape of all possible amino acid substitutions at every position; we then defined highly variable (hotspot) regions as contiguous sequence segments in which substitutions display high variance in editing efficiency, contain multiple variants that significantly surpass wild-type efficiency (above a predefined threshold, e.g. > X % increase), and overlap or lie in close proximity (within a few residues) to the nucleic acid-proximal or PAM-proximal residues defined above. These hotspot regions were prioritized for subsequent engineering of LaTranC variants.

### Construction of mutant libraries

For the TranC amino acid sites identified during site selection, saturation mutagenesis was performed by introducing an NNK codon at each target position using areverse PCR (Beijing Tsingke Biotech). The amino acid site screened by P1a and P1b is provided in **Supplementary Table 1**). As an example, for amino acid positions 76 to 80 of eLaTranC, the forward primer (CTAGCTACAAGAAGATG**NNKAATAAGACCTGG**CGGAAGGCTCTGATCGACC) was designed to incorporate the NNK mutation at position 76, while the downstream primer (CATCTTCTTGTAGCTAGGCCACAC) was used to amplify the P1 plasmid. Similarly, separate upstream primers containing NNK mutations (indicated in bold) were designed for positions 77 to 80, paired with the same downstream primer for amplification. The five groups of PCR products were analyzed by agarose gel electrophoresis, and the expected bands were excised and purified using a gel extraction kit (Thermo Fisher Scientific). The purified products were then digested with DpnI restriction endonuclease (Vazyme Biotech) at 37 °C for 2 hours, followed by enzyme inactivation at 80 °C for 20 minutes. The digested fragments were subsequently ligated using a seamless cloning kit (GeneSand). The five ligation products were combined and purified with a DNA purification kit (Sangon). 100 ng of the purified ligation product (A typical transformation using 5 to 8 mixed products amplifying by the same pair of primer sets) was electroporated (Lonza Biosciences) into DH10B electrocompetent cells (WEIDI) with program GE 100, then cells were immediately resuspended in 1 mL SOC medium and recovered at 37 °C with shaking at 220 rpm for 10 minutes. The transformed culture was spread onto LB agar plates supplemented with ampicillin and incubated overnight at 37 °C. Positive clones were scraped and plasmid DNA was extracted using *EasyPure*® HiPure Plasmid MiniPrep Kit (TransGene Biotech).

### Screening condition test

Initially, the P2 plasmid expressing the *ccdB* gene (constructed from p11-LacY-wtx1^20^) was introduced into BW25141 cells and used to prepare electrocompetent cells. BW25141 cells harboring P2 plasmid were then electrical transformed with either 100 ng P1a-eLaTranC or P1a-deadTranC (D253A) plasmids by 4D-Nucleofector (Lonza Biosciences) with program GE 100. Cells after electrocuted were recovered in SOC medium at 37°C with shaking at 220 rpm for either 10 or 30 minutes. Then recovered cultures were plated on LB agar plates supplemented with ampicillin and different concentrations of arabinose (0, 4, or 10 mM; Solarbio) and incubated overnight at 37°C. Colony formation was quantified to assess cell survival (**Supplementary Table 2**).

### Library screening

The constructed subpool plasmids were mixed according to the amino acid positions. For example, all mutant plasmid libraries with amino acid residues at positions 61-120 of eLaTranC were mixed in equal amounts. 100 ng of mixed plasmid library were electrical transformed into BW25141 cells harboring P2 plasmid by 4D-Nucleofector (Lonza Biosciences) with program GE 100. Cells after electric shock recovered in 1 mL SOC medium at 37°C with shaking at 220 rpm (10 min for P1a; 20 min for P1b). The recovered cultures were plated on LB agar plates supplemented with ampicillin and L-arabinose (P1a: 8 mM; P1b: 12 mM) and incubated overnight at 37°C. Surviving colonies were subjected to sequencing. Under the P1a selection condition, more than 50,000 colonies were obtained, from which 71 clones were randomly selected for Sangar sequencing. Under the P1b condition, all 1,058 surviving colonies were sequenced by high-throughput sequencing. All the amino acid mutations that were screened and successfully sequenced are provided in (**Supplementary Table 1**).

### Plant protoplasts transfection

Plant protoplast transformation was performed as described previously^21^ with modifications. Briefly, aerial tissues from 10–14-day-old maize seedlings (6WC) were dissected into 0.5–1 mm strips and digested in an enzyme solution (1.5% Cellulase R10, 0.75% Macerozyme R10, 0.6 M mannitol, 10 mM MES pH 5.7, 10 mM CaCl_2_, 0.1% BSA) in the dark at 25 °C with gentle shaking (40–60 r.p.m.) for 4–6 h (maize). The digestate was filtered through a 100-μm mesh, and the filtrate was diluted 1:1 with W5 solution (154 mM NaCl, 125 mM CaCl_2_, 5 mM KCl, 5 mM glucose, 2 mM MES pH 5.7) and centrifuged at 100 g for 10 min. The pellet was resuspended in ice-cold W5 and incubated on ice for 30 min. After centrifugation, protoplasts were resuspended in MMg solution (0.6 M mannitol, 15 mM MgCl_2_, 4 mM MES pH 5.7) and counted. Cells were adjusted to 2 × 10^5^ cells/mL and kept on ice. For transformation, 5 μg editor DNA and 5 μg sgRNA plasmid were added to 100 μL protoplast suspension, followed by 110 μL PEG/Ca^2+^ solution (40% PEG4000, 0.6 M mannitol, 100 mM CaCl_2_). After gentle mixing and 30 min incubation at room temperature, the reaction was quenched with 880 μL W5 solution. Protoplasts were pelleted (100 g, 3 min), resuspended in 1 mL WI solution (0.6 M mannitol, 4 mM KCl, 20 mM MES pH 5.7). Transfected protoplasts were cultured in the dark at 25 °C for 48 h before analysis. After incubation, the protoplasts were collected, and genomic DNAs were extracted for deep amplicon sequencing.

## Acknowledgements

This work was supported by the Agriculture Science and Technology Major Project, the National Key Research and Development Program (2022YFF1002802), the National Natural Science Foundation of China (32230088, 32250012, and 32201224), the CAS Projects for Young Scientists in Basic Research (YSBR-080), and the New Cornerstone Science Foundation. The figure schematics in Figure 4 were made using BioRender.

## Contributions

K.T.Z. and Z.Z. conceptualized the study. Quan Gao and Z.Z. assembled the vectors and conducted experiments in HEK293T cells. L.L. conducted protoplast transfection experiments. M.H., Quan Gao, and L.Z. performed amplicon sequencing. Z.Z., Quan Gao, and S.L. collected and analyzed amplicon sequencing data. Qiang Gao and H.J. performed the AI Training. Qiang Gao and Z.W. wrote scripts and processed the raw amplicon sequencing data. S.J. and C.G. led the discovery of eLaTranC and advised on engineering. Z.Z. and K.T.Z. prepared the figures. Z.Z. and K.T.Z. wrote the manuscript with input from all authors.

